# Genetic diversity and population structure of cowpea [*Vigna unguiculata* (L.) Walp.] accessions from Togo using SSR Markers

**DOI:** 10.1101/2021.05.17.444438

**Authors:** Yao Dodzi Dagnon, Koffi Kibalou Palanga, Damigou Bammite, Ghislain Comlan Akabassi, Koffi Tozo

## Abstract

Cowpea [*Vigna unguiculata* (L.) Walp.] is a crop with significant agronomic and nutritional potential. I is very appreciate by local people. It is the third food habit in Togo after maize and rice. However, several accessions of cowpea cultivated in Togo are now prone to extinction, creating a risk of genetic erosion. It is therefore urgent to assess the genetic diversity of accessions in order to set up a good conservation program. To achieve this, genetic diversity and phylogenetic relationships among 70 accessions of cowpea collected in the five (5) administrative regions of Togo were assessed using Simple Sequence Repeat (SSR) molecular markers. Twenty-eight out of the thirty-two (32) primer pairs screened for polymorphism were polymorphic, and a total of 164 alleles were detected for the 28 loci with an average of 5.82 alleles per locus. Polymorphic Information Content (PIC) values ranged from 0.18 to 0.895, with an average value of 0.67. Population structure analysis using model-based revealed that the cowpea germplasm was grouped into two subpopulations. The analysis of molecular variance (AMOVA) revealed that 85% of genetic variation existed among individuals within regions. The fixation index (Fst) value, which was 0.018, was low, indicating relatively low population differentiation. The Togolese cowpea germplasm collection was grouped into four groups independently of theirs origins. This study provides a foundation for a Togolese cowpea germplasm conservation program and can serve for the selection of parental material for further studies aimed at the genetic improvement of local germplasm.

## Introduction

Cowpea [*Vigna unguiculata* (L.) Walp.] is an important food legume in developing countries of the tropics and subtropics, especially in Sub-Saharan Africa, Asia, and Central and South America [1,2], and in some temperature area, including the Mediterranean region and the southern states of the USA [3,4]. Its global annual production is 3.5 million metric tons, and Nigeria alone produced over 2.24 million metric tons on 2.52 million ha, followed by Niger, which produced 1.77 million metric tons on 5.57 million ha in 2017 [5]. Cowpea is commonly cultivated as a nutritious and highly palatable food source. The seed is reported to contain 24% crude protein, 53% carbohydrates, and 2% fat [6]. Referred to as the ‘poor man’s meat’ because of its good protein quality and high nutritional value [7], cowpea hay is also useful in the feeding of animals during the dry season in many parts of West Africa [8,9]. All parts of the cowpea are used for food. The leaves, green pods, green peas and dry grains are consumed as different dishes. Cowpea plays a very important subsistence role in the diets of many households in Africa [10]. It also has an economic value to the farming households since it is also a cash crop [11]. Besides, cowpea is a valuable component of farming systems in many areas because of its ability to restore soil fertility through nitrogen fixation for succeeding cereal crop grown in rotation with it [8,12,13].

Nowadays, with problems of climatic variability and increasing world population, the demand for food also increases. Unfortunately, in Togo, many cowpea landraces are abandoned for their seed color or various reasons [6]. The climatic variability leads farmer to select the landraces which have a short vegetative cycle. All those facts lead to genetic erosion of the crop [2]. A major goal of cowpea breeding and genetic improvement programs around the world is to combine desirable agronomic traits (e.g. time to maturity, photoperiod sensitivity, plant type and seed quality) with resistance to the major biotic stresses threatening the crop production [9,14].

Genetic diversity is the extent to which heritable material differs within a group of plants and results from evolution, including domestication and plant breeding. Assessing the genetic diversity of cowpea germplasm is a prerequisite for effective breeding and germplasm conservation. Genetic studies of cowpea diversity have been done in several countries using DNA molecular markers such as random amplified polymorphic DNA (RAPD) [7,15,16], amplified fragment length polymorphisms (AFLP) [17], restriction fragment length polymorphisms (RFLP) [18], inter-simple sequence repeat (ISSR) [4] and simple sequence repeat (SSR) [9,19]. Of all these markers, SSR is the most widely used marker in genetic diversity analysis due to their multiallelic nature, high reproducibility, co-dominant inheritance, abundance and extensive genome coverage that has already been reported for crops like pigeon pea [20,21] or rice [22,23,24,25]. In cowpea, the earliest use of SSR for assessing the crop genetic diversity was conducted by Li et al.[19]. SSRs are also used to identify genotype, seed purity evaluation and variety protection, pedigree analysis and genetic mapping of simple and quantitative traits and marker-assisted selection breeding [9,19,26]. Prior to this study, there was no study conducted on genetic diversity of cowpea germplasm. The present study was, therefore, undertaken to address the knowledge gap on the country germplasm genetic diversity based on SSR markers.

## Materials and methods

### Plant materials

The plant material consists of 70 accessions of cowpea [*Vigna unguiculata* (L.) Walp.] collected from producers in the five (05) regions of Togo (Maritime, Plateaux, Centrale, Kra and Savannah) between 2014 and 2016 (Table 1). Among the 70 cowpea accessions, three are 3 varieties listed in the national catalogue of species and varieties grown in Togo. These varieties are VITOCO and TVX bred by IITA-IBADAN, and VITA5 bred by the University of Ifê (Nigeria) and are widely cultivated in Togo. They were obtained from the Togolese Institute of Agriculture Research (ITRA). The 70 cowpea accessions represent the collection from all major growing areas of cowpea in Togo. Their local name and place of collection are provided in Table 1. In our study, all accessions from a region represented a population.

**Table 1.** List of the Cowpea accessions, characteristics and collection.

| Accessions | Growth habit | Flower color | Seed size | Seed color | Status | Regions |
| --- | --- | --- | --- | --- | --- | --- |
| Yélengo | Creeping | Purple | small | Beige red | Landrace | Centrale |
| Gbedéfouba | Creeping | Purple | small | Beige red | Landrace | Centrale |
| Guinsibibè | Creeping | White | big | white | Landrace | Centrale |
| Sotouboua | Creeping | White | big | white | Landrace | Centrale |
| Hèkou hèkou | Creeping | White | medium | white | Landrace | Centrale |
| Tchéwo | Creeping | White | small | white | Landrace | Centrale |
| Tchéwo koumoka | Creeping | White | small | white | Landrace | Centrale |
| Vitoco 2 | Erected | Purple | small | white | Breeding Line | Centrale |
| Komi | Creeping | White | small | white | Landrace | Centrale |
| Vita 5 | Creeping | White | small | white | Breeding Line | Centrale |
| Kétchéyi soukpèlo | Erected | Purple | small | purple red | Landrace | Kara |
| Kétchéyi Koussémo | Creeping | Purple | small | Red wine | Landrace | Kara |
| Kétchéyi | Erected | Purple | small | Burgundy purple | Landrace | Kara |
| Djodjowou | Creeping | White | big | white | Landrace | Kara |
| Koufaldo | Creeping | White | big | white | Landrace | Kara |
| Dapango kaga | Creeping | White | medium | white | Landrace | Kara |
| Dapango Koukpèto | Creeping | White | medium | white | Landrace | Kara |
| Kandjarga | Creeping | White | medium | Yellow sand | Landrace | Kara |
| Lamga | Creeping | White | small | white | Landrace | Kara |
| Simpayo | Creeping | White | small | white | Landrace | Kara |
| Tinkou | Creeping | White | small | white | Landrace | Kara |
| Sodjadéawoudadè | Semi erected | Purple | medium | Beige red | Landrace | Maritime |
| Togbéyi | Creeping | Purple | medium | Beige red | Landrace | Maritime |
| Amélassiwa | Semi erected | White | small | white | Landrace | Maritime |
| Dakarvi | Creeping | White | small | white | Landrace | Maritime |
| Kpédéviyi | Creeping | Purple | small | Beige red | Landrace | Maritime |
| Kpédévi | Creeping | Purple | small | Beige red | Landrace | Maritime |
| Téklikoé | Creeping | Purple | small | Rouge noir | Landrace | Maritime |
| Damadoami | Creeping | Purple | small | purple red | Landrace | Maritime |
| Itouloka | Creeping | Purple | small | Burgundy purple | Landrace | Maritime |
| Assiamaton | Semi erected | White | medium | white | Landrace | Maritime |
| Yéboua | Creeping | White | medium | white | Landrace | Maritime |
| Agnokoko | Creeping | White | small | white | Landrace | Maritime |
| Amélassiwa 2 | Creeping | White | small | white | Landrace | Maritime |
| Gban molou | Creeping | White | small | white | Landrace | Maritime |
| Sakawouga | Creeping | Purple | medium | Reddish grey | Landrace | Plateaux |
| Ayi djin | Erected | Purple | medium | Beige red | Landrace | Plateaux |
| Tcharabaou djin | Creeping | Purple | medium | Red wine | Landrace | Plateaux |
| TVX | Erected | White | small | white | Breeding Line | Plateaux |
| Poli poli | Creeping | Purple | small | Beige red | Landrace | Plateaux |
| 45 jours rouges | Erected | Purple | small | purple red | Landrace | Plateaux |
| Maca | Creeping | Purple | small | Red wine | Landrace | Plateaux |
| Kéchéyi 2 | Semi erected | Purple | small | Burgundy purple | Landrace | Plateaux |
| Azangba | Erected | Purple | small | Burgundy purple | Landrace | Plateaux |
| Agamassikè | Creeping | White | medium | white | Landrace | Plateaux |
| Amélassiwa 3 | Creeping | White | medium | white | Landrace | Plateaux |
| Sotoco | Creeping | White | medium | white | Landrace | Plateaux |
| Vitoco | Semi erected | White | medium | white | Breeding Line | Plateaux |
| Atakpamé | Creeping | White | medium | white | Landrace | Plateaux |
| Pamplovi | Creeping | White | small | white | Landrace | Plateaux |
| Siéloune | Semi erected | White | small | Yellow Gold | Landrace | Savannah |
| Malgbong bomoine | Semi erected | Purple | small | purple red | Landrace | Savannah |
| Esatoune | Creeping | Purple | small | Red wine | Landrace | Savannah |
| Simporé | Creeping | White | big | white | Landrace | Savannah |
| Atougbanda | Semi erected | White | medium | white | Landrace | Savannah |
| Bieng nomio | Creeping | White | medium | white | Landrace | Savannah |
| Golenga | Creeping | White | medium | white | Landrace | Savannah |
| Malgbong bopiel | Creeping | White | medium | white | Landrace | Savannah |
| Pélam | Creeping | White | medium | white | Landrace | Savannah |
| Alacante | Semi erected | White | medium | white | Landrace | Savannah |
| Toi | Semi erected | White | medium | white | Landrace | Savannah |
| Bieng oune | Creeping | White | small | white | Landrace | Savannah |
| Etougnognoli | Creeping | Purple | small | white | Landrace | Savannah |
| Etoukakali | Creeping | White | small | white | Landrace | Savannah |
| Gouarga | Creeping | White | small | white | Landrace | Savannah |
| Itouloka | Creeping | White | small | white | Landrace | Savannah |
| Kampirigbène | Semi erected | White | small | white | Landrace | Savannah |
| Natoguildjole | Creeping | White | small | white | Landrace | Savannah |
| Téléga | Semi erected | White | small | white | Landrace | Savannah |
| Toboni | Creeping | White | small | white | Landrace | Savannah |

### DNA extraction

The genomic DNA was extracted from fresh leaf material of 21 day-old-plants of each of the 70 cowpea accessions following the mixed alkyl trimethyl ammonium bromide (MATAB) protocol described by Risterucci et al. [27]. The quality of the extracted DNA was checked on 0.8% agarose gel, and its concentration was estimated by comparing the obtained bands with the bands of a Smart Ladder (MW-1700-10-Eurogentec) of known concentration. The working DNA concentration was then adjusted to 25 ng/μL. The DNA samples were analysed at the Centre d’Etude Régional pour l’Amélioration de l’Adaptation à la Sécheresse (CERAAS) in Senegal.

### Polymerase Chain Reaction using SSR markers

A total of 28 polymorphic SSR markers was used to screen 70 cowpea DNA samples (Table 2). The forward and reverse primers of each of the 28 SSR markers (Table 2) were labeled at their 5’ end with fluorescent dyes to enable detection. The PCR reaction was conducted in a total volume of 10 μl, (5 μl of DNA and 5 μl of a PCR solution). The PCR solution was prepared with 55 μL of 10X buffer, 55 μL of dNTPs (200 μg), 22 μL of MgCl_2_ (0.5 mM), 9 μL of each primer (0.1 μM), 9 μL of IR dye (0.1 μM), 55 μL of Taq Polymerase 1U and 227 μL of ultrapure water. The PCR reaction was carried out in a 96-block thermal cycler (MWG AG biotech). The thermal cycling conditions were as follows: initial denaturation step at 94 °C for 4 min followed by 26 cycles of denaturation (94 °C) for 60 s, hybridization (50 - 55 °C according to the primers) for 1 min, primer extension (72°C) for 1 min 15 seconds, followed by a final extension at 72 °C for 7 min. After PCR, a 0.8% agarose gel was used to control the quality of the amplification products. The PCR plates were then covered with aluminium foil to prevent fluorochrome degradation and placed in a refrigerator.

**Table 2.**
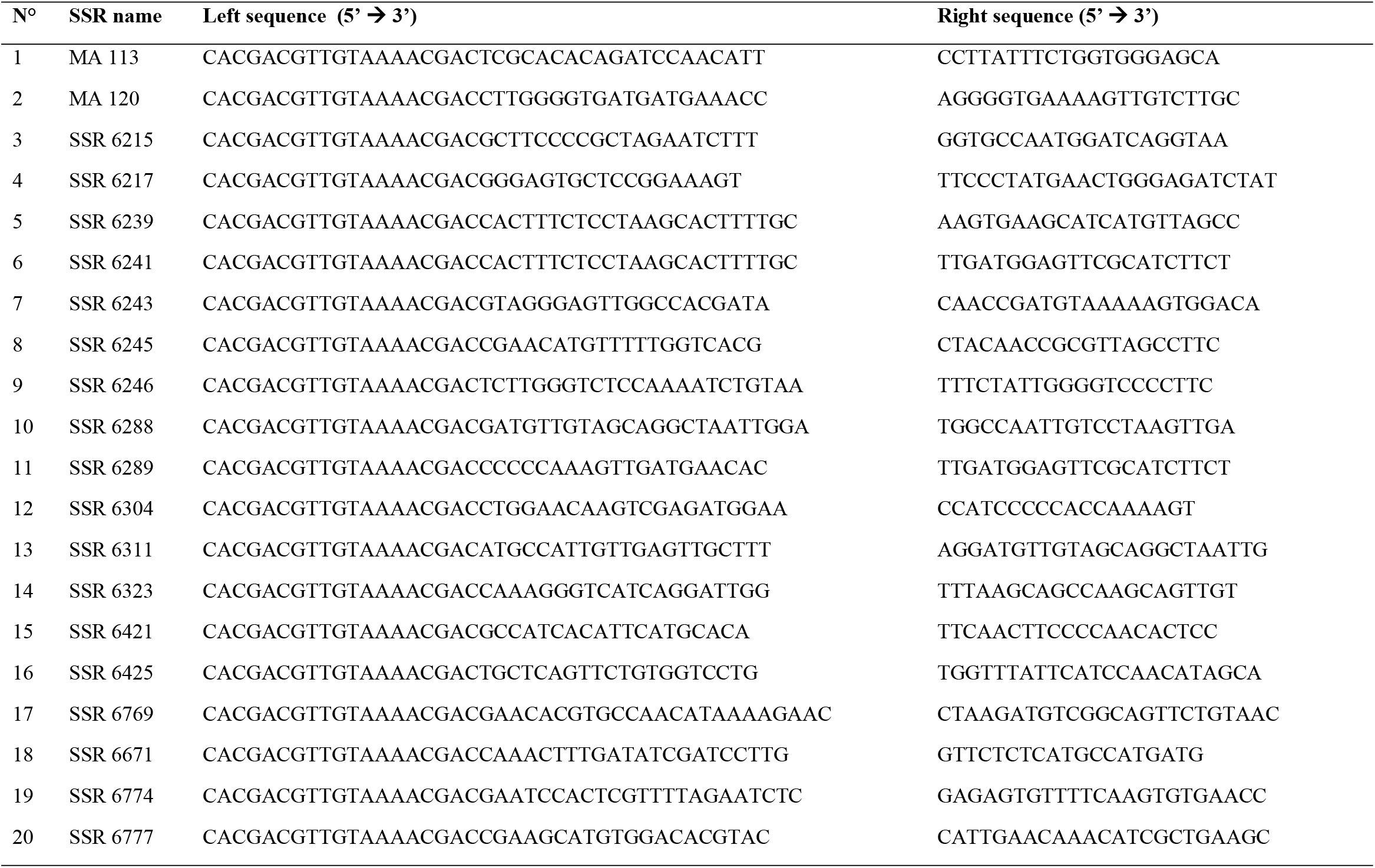
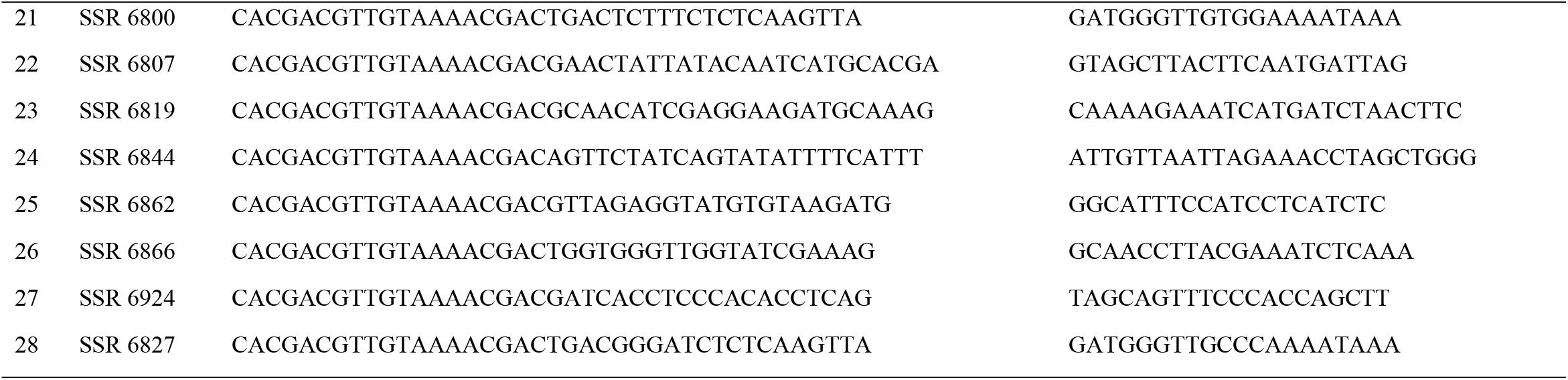
Primer sequences of the 28 simple sequence repeat (SSR) markers used in this study [33].

### Gel electrophoresis

The amplification PCR products were analyzed by electrophoresis on a 6.5% polyacrylamide denaturing gel on Licor 4300 sequencer (LICOR Inc., NE, USA). Before loading the gel, the multiplexed PCR products were denatured at 94 ° C for 3 min, and then the plate was placed on ice. The amount of denatured DNA loaded in the wells of the deposition rack was 2.5 μL. An infrared camera detected the fluorescence signals emitted by the marked fragments when excited with laser diodes at two different wavelengths (682 and 782 nm). The images were automatically recorded and downloaded for analysis. Allele sizes were estimated by comparing with different bands of the size marker (ladder produced by CIRAD).

### Scoring of bands and data analysis

All images of the gel profiles were printed for reading. A binary matrix was generated for all accessions based on the patterns of the bands observed at a particular locus. The GenAlex 6.4 software [28] was used to determine genetic parameters such as the total number of alleles per locus, major allele frequency, observed heterozygosity, expected heterozygosity and the polymorphism information content (PIC) values for each SSR locus. An analysis of molecular variance (AMOVA) was performed to test the degree of differentiation among and within the sources of collection of the cowpea accessions. Finally, the software DARwinV.5.0.158 [29] was used to make the dendrogram.

The population structure of the 70 cowpea accessions was established using the Bayesian clustering method in STRUCTURE version 2.3.2 [30]. The length of the burn-in period and Markov Chain Monte Carlo (MCMC) were set at 10,000 iterations [31]. To obtain an accurate estimation of the number of populations, ten runs for each K-value were performed with K ranging from 1 to 10. Further, Delta K values were calculated, and the appropriate K value was estimated by implementing the method of Evanno et al. [31] using the STRUCTURE Harvester program [32].

## Results

### Genetic Polymorphism SSR Markers

Out of 32 SSR primer pairs tested, only 28 generated clear profile and were polymorphic. A total of 164 alleles were generated by those 28 markers across the 70 accessions. The number of alleles detected per SSR primer pairs varied between two (2) to fourteen (14), with an average of 5.86 alleles per loci. The lowest number of alleles per locus was detected for the markers SSR6217, SSR6774, SSR6311, SSR6243, SSR6671, and SSR6288. The highest number of alleles was recorded for SSR6800. A total of 18 rare alleles were detected in this study. The number of effective alleles per marker ranged from 1.21 to 6.44, with an average of 3.05 with the markers SSR6571 and the marker SSR6807 having respectively the lowest and the highest number of effective alleles respectively. For the SSR loci, polymorphism information content (PIC) representing a measure of the allelic diversity for a specific locus varied from 0.20 to 0.89 with an average of 0.58. Ten SSR loci (SSR6243, SSR6215, SSR6819, SSR6800, SSR6239, SSR6807, SSR6844, MA120, SSR6866 and MA113) exhibited PIC values higher than 0.70, indicating their usefulness in discriminating genotypes. The observed heterozygosity values ranged from 0.0 to 0.35 with an average of 0.07, and the major allele frequency varied from 16.17% to 89.06%. This study has also revealed a number of rare alleles and unique alleles. The rare alleles represented near 11% of the whole detected alleles, with a total of 14 rare alleles and a total of 3 unique alleles were also detected. SSR6800, SSR6245 and SS6215 have respectively produced an unique allele for the accession Amélassiwa 3, Kampirigbène and Agnokoko (Table 3).

**Table 3.** The number of alleles per locus, Major allele frequency, Expected heterozygosity, Observed heterozygosity and polymorphism information content (PIC) of the 28 SSR markers across 70 cowpea accessions.

| Markers | Alleles |  |  | Number of effective alleles<sup>b</sup> | Major allele frequency (%)<sup>c</sup> | Expected heterozygosity<sup>d</sup> | Observed heterozygosity<sup>e</sup> | PIC<sup>f</sup> |
| --- | --- | --- | --- | --- | --- | --- | --- | --- |
|  | Number<sup>a</sup> per loci | Unique | Rare |  |  |  |  |  |
| SSR6421 | 3 | 0 | 1 | 1.99 | 55.70 | 0.43 | 0.00 | 0.50 |
| SSR6246 | 3 | 0 | 1 | 1.45 | 81.20 | 0.28 | 0.01 | 0.31 |
| SSR6217 | 2 | 0 | 0 | 1.86 | 63.60 | 0.43 | 0.02 | 0.46 |
| SSR6323 | 3 | 0 | 0 | 2.10 | 57.10 | 0.46 | 0.02 | 0.52 |
| SSR6769 | 6 | 0 | 0 | 4.95 | 26.07 | 0.65 | 0.02 | 0.80 |
| SSR6425 | 4 | 0 | 0 | 2.36 | 57.73 | 0.56 | 0.02 | 0.58 |
| SSR6774 | 2 | 0 | 0 | 1.87 | 63.30 | 0.46 | 0.01 | 0.46 |
| SSR6777 | 3 | 0 | 0 | 1.70 | 73.71 | 0.41 | 0.01 | 0.36 |
| SSR6311 | 2 | 0 | 0 | 1.94 | 59.00 | 0.47 | 0.08 | 0.48 |
| SSR6862 | 5 | 0 | 0 | 2.68 | 56.41 | 0.56 | 0.02 | 0.63 |
| SSR6243 | 2 | 0 | 0 | 1.69 | 71.29 | 0.38 | 0.03 | 0.41 |
| SSR6215 | 11 | 1 | 1 | 5.91 | 30.45 | 0.79 | 0.34 | 0.83 |
| SSR6924 | 4 | 0 | 1 | 1.91 | 68.74 | 0.46 | 0.01 | 0.48 |
| SSR6671 | 2 | 0 | 0 | 1.26 | 88.48 | 0.20 | 0.03 | 0.20 |
|  | Number<sup>a</sup> per loci | Unique | Rare |  |  |  |  |  |
| SSR6819 | 10 | 0 | 1 | 6.66 | 23.37 | 0.75 | 0.00 | 0.85 |
| SSR6800 | 14 | 1 | 3 | 8.17 | 21.34 | 0.83 | 0.01 | 0.88 |
| SSR6245 | 3 | 1 | 0 | 1.70 | 71.29 | 0.38 | 0.05 | 0.41 |
| SSR6304 | 4 | 0 | 1 | 2.09 | 59.51 | 0.49 | 0.03 | 0.52 |
| SSR6288 | 2 | 0 | 0 | 1.52 | 77.94 | 0.34 | 0.38 | 0.34 |
| SSR6239 | 9 | 0 | 0 | 6.04 | 24.03 | 0.78 | 0.02 | 0.83 |
| SSR6807 | 13 | 0 | 1 | 9.21 | 16.17 | 0.83 | 0.08 | 0.89 |
| SSR6241 | 7 | 0 | 0 | 2.47 | 61.20 | 0.58 | 0.32 | 0.59 |
| SSR6844 | 11 | 0 | 2 | 6.69 | 25.76 | 0.81 | 0.02 | 0.85 |
| MA120 | 10 | 0 | 1 | 5.58 | 26.65 | 0.77 | 0.03 | 0.82 |
| SSR6866 | 11 | 0 | 0 | 7.14 | 28.42 | 0.80 | 0.00 | 0.86 |
| MA113 | 12 | 0 | 3 | 7.83 | 23.71 | 0.79 | 0.00 | 0.87 |
| SSR6289 | 3 | 0 | 1 | 1.25 | 89.06 | 0.19 | 0.19 | 0.20 |
| SSR6827 | 3 | 0 | 1 | 1.63 | 74.69 | 0.38 | 0.27 | 0.39 |

<sup>a</sup> Total (164), Average (5.86), Minimum (2), Maximum (14); <sup>b</sup> Average (3.63), Minimum (1.25), Maximum (9.21); <sup>c</sup> Average (52.71), Minimum (16.17), Maximum (89.06); <sup>d</sup> Average (0.54), Minimum (0.19), Maximum (0.83); <sup>e</sup> Average (0.07), Minimum (0.00), Maximum (0.38); <sup>f</sup> Average (0.58), Minimum (0.20), Maximum (0.89).

### Genetic relationship of cowpea populations

SSR markers used in this study revealed high percentages of polymorphic loci (average of 99.28%). The lowest percentage of polymorphism was observed for cowpea population one, corresponding to the Centrale region, while the percentage of polymorphism observed for each of the other four regions was 100%. The number of alleles detected in each population is not uniform and varied from 102 alleles in the cowpea population from the Centrale region to 127 alleles in the population of the Plateaux region. Among the five population investigated, the mean values of observed alleles (Na) and effective alleles (Ne) were 3.96 and 2.92, respectively. Population 3 (from Centrale Region) recorded the lowest value of Na (3.64), while the highest value (4.54) was recorded from population 5 (from Savane Region). For the effective number of alleles, the lowest value (2.85) was recorded from population 3 (from Maritime Region), while the highest value (3.07) was displayed by population 5. and Ne (2.10). Shannon’s information index ranged from 1 to 1.08 with a mean of 1.03. The observed heterozygosity (Ho) ranged from 0.07 (population 5, population 4 and population 3) to 0.08 (population 1 and population 2) with an average of 0.07. The expected heterozygosity (He) was moderately high and ranged from 0.53 (Population 2) to 0.55 (Population 1 and Population 3), with an average of 0.54. The unbiased expected heterozygosity ranged from 0.56 (Population 2, Population 4 and Population 5) to 0.58 (Population 1 and Population 3), with an average of 0.46. According to the results, the five regions displayed almost similar diversity of cowpea. The values for the inbreeding coefficient expressed by the fixation index F ranged from 0.79 (Population 5) to 0.85 (Population 1 and Population 3) with an average of 0.82 at the population level (Table 4). Genetic similarity among the five populations was high and ranged from 0.85 between Population 1 and Population 4 to 0.94 between Population 3 and Population 4 (Table 5).

**Table 4.** Summary of different cowpea population diversity statistics averaged over the 28 SSR loci.

| Population | Size | Allele number | % P | Na | Ne | I | Ho | He | uHe | F |
| --- | --- | --- | --- | --- | --- | --- | --- | --- | --- | --- |
| **Centrale** | 10 | 102 | 96.43 | 3.64 | 2.86 | 1.015 | 0.080 | 0.550 | 0.580 | 0.846 |
| **Kara** | 11 | 105 | 100 | 3.75 | 2.86 | 0.997 | 0.083 | 0.529 | 0.556 | 0.821 |
| **Maritime** | 14 | 111 | 100 | 3.96 | 2.84 | 1.046 | 0.068 | 0.555 | 0.577 | 0.854 |
| **Plateaux** | 15 | 110 | 100 | 3.93 | 2.94 | 1.031 | 0.066 | 0.543 | 0.563 | 0.806 |
| **Savannah** | 20 | 127 | 100 | 4.54 | 3.07 | 1.077 | 0.067 | 0.541 | 0.555 | 0.788 |
| **Mean** | **14** | **111** | 99.28 | 3.96 | 2.92 | 1.033 | 0.073 | 0.544 | 0.566 | 0.823 |
| **SE** | **1.16** | **4.32** |  | 0.19 | 0.15 | 0.047 | 0.012 | 0.018 | 0.019 | 0.028 |

Population 1 =cowpea accessions from Centrale Region, Population 2 = cowpea accessions from Kara Region, Population 3 = cowpea accessions from Maritime Region, Population 4 = cowpea accessions from Plateaux Region, Population 5: cowpea accessions from Savannah Region Na = Number of different alleles; Ne = number of effective alleles, I = Shannon's Information Index, Ho = observed heterozygosity, He = expected heterozygosity, uHe = Unbiased expected heterozygosity, F = Fixation index.

**Table 5.** Genetic similarities between cowpea populations.

| Population 1 | Population 2 | Population 3 | Population 4 | Population 5 |  |
| --- | --- | --- | --- | --- | --- |
| 1.000 |  |  |  |  | **Population 1** |
| 0.898 | 1 |  |  |  | **Population 2** |
| 0.857 | 0.900 | 1 |  |  | **Population 3** |
| 0.847 | 0.865 | 0.945 | 1 |  | **Population 4** |
| 0.849 | 0.874 | 0.901 | 0.922 | 1 | **Population 5** |

The genetic differentiation indices between populations (Fst) varied from 0.00 (between Centrale and Kara Regions, Kara and Savannah Regions, Maritime and Plateaux Regions) to 0.057 between Centrale and Maritime Regions. Differentiation appears to be null or low between accessions from different regions except for Centrale and Maritime Regions, which appears moderate (Table 6).

**Table 6:** Pairwise Fst values of the accessions.

| Centrale | Kara | Maritime | Plateaux | Savannah |  |
| --- | --- | --- | --- | --- | --- |
| 0.000 |  |  |  |  | **Centrale** |
| 0.000 | 0.000 |  |  |  | **Kara** |
| 0.057 | 0.030 | 0.000 |  |  | **Maritime** |
| 0.024 | 0.010 | 0.000 | 0.000 |  | **Plateaux** |
| 0.012 | 0.000 | 0.039 | 0.028 | 0.000 | **Savannah** |

### Population Structure of the 70 cowpea accessions based on 28 SSR markers

The analysis of the population structure based on the ΔK value grouped the seventy accessions into two subpopulations (Fig 2). Membership of all genotypes to a particular subpopulation was based on a likelihood threshold of 0.55. Subpopulation 1 had the largest membership with 64.28% of the accessions, while the smallest Cluster 1 only gathered 35.71% of the accessions (Table 7). Based on the threshold of 0.55, the study did not reveal any admixture among the accessions. Both subpopulations were composed of accessions from the five regions, and white-colored seeds dominated both. The first subpopulation comprised five accessions from the Centrale region, seven accessions from Kara region, nine accessions from Maritime region, 11 accessions from the Plateaux region and 13 accessions of the Savane region while subpopulation comprises five accessions from the Centrale region, fourth accessions from the Kara region, five accessions from the Maritime region, fourth accessions Plateaux region and 7 accessions of Savane region. Subpopulation 1 had more accessions from Savane and Plateaux regions, while population 2 had almost equal access to each region. Regarding the seeds coat color, subpopulation 1 was the most heterogeneous and included 66.67% of white-colored seeds, 11.11% of beige red-colored seeds, 8.89% of red wine-colored seeds and 4.44% of burgundy purple-colored seeds, while subpopulation 2 included 64% of white-colored seeds, 12% of beige red-colored seeds, 12% of purple -colored seeds and 8% of burgundy purple-colored of the reddish grey, golden yellow, blackish red and purple red colors were recorded respectively for one genotype in subpopulation one while the genotype presenting those seed color were absent in subpopulation 2.

**Table 7.**
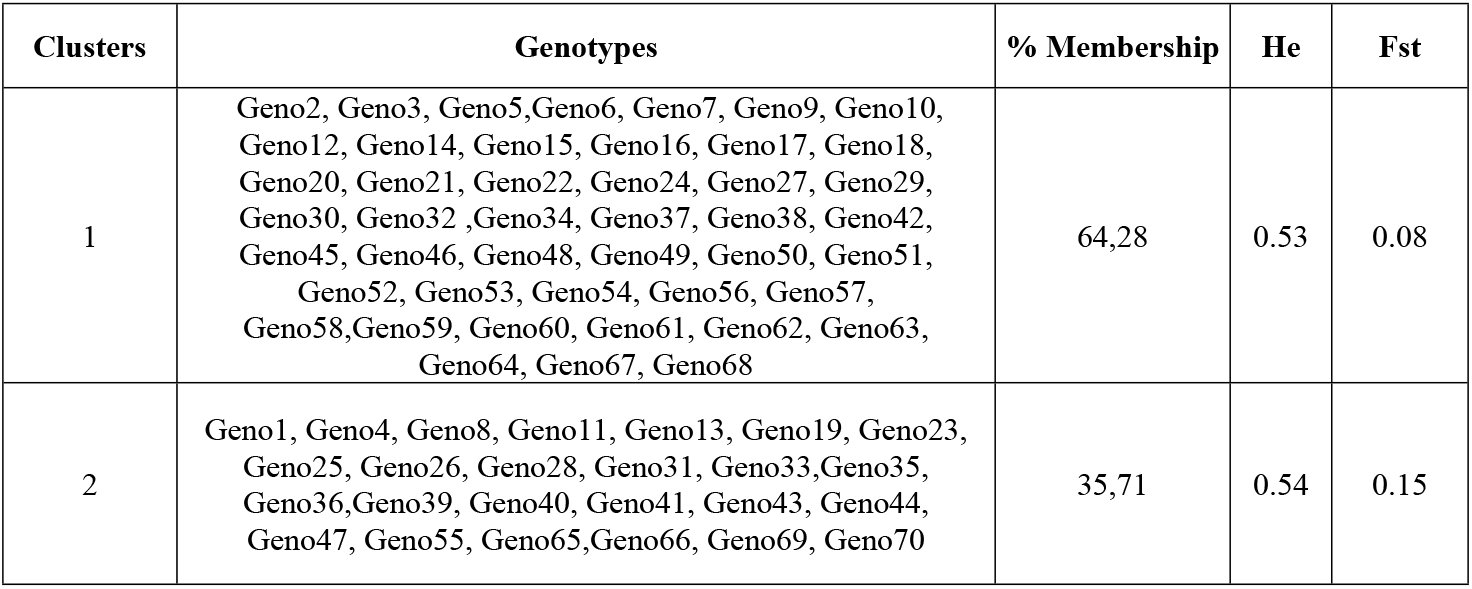
Analysis of molecular variance (AMOVA) based on 28 SSR markers.

| Clusters | Genotypes | % Membership | He | Fst |
| --- | --- | --- | --- | --- |
| 1 | Geno2, Geno3, Geno5, Geno6, Geno7, Geno9, Geno10, Geno12, Geno14, Geno15, Geno16, Geno17, Geno18, Geno20, Geno21, Geno22, Geno24, Geno27, Geno29, Geno30, Geno32, Geno34, Geno37, Geno38, Geno42, Geno45, Geno46, Geno48, Geno49, Geno50, Geno51, Geno52, Geno53, Geno54, Geno56, Geno57, Geno58, Geno59, Geno60, Geno61, Geno62, Geno63, Geno64, Geno67, Geno68 | 64,28 | 0.53 | 0.08 |
| 2 | Geno1, Geno4, Geno8, Geno11, Geno13, Geno19, Geno23, Geno25, Geno26, Geno28, Geno31, Geno33, Geno35, Geno36, Geno39, Geno40, Geno41, Geno43, Geno44, Geno47, Geno55, Geno65, Geno66, Geno69, Geno70 | 35,71 | 0.54 | 0.15 |

**Table 7a:**
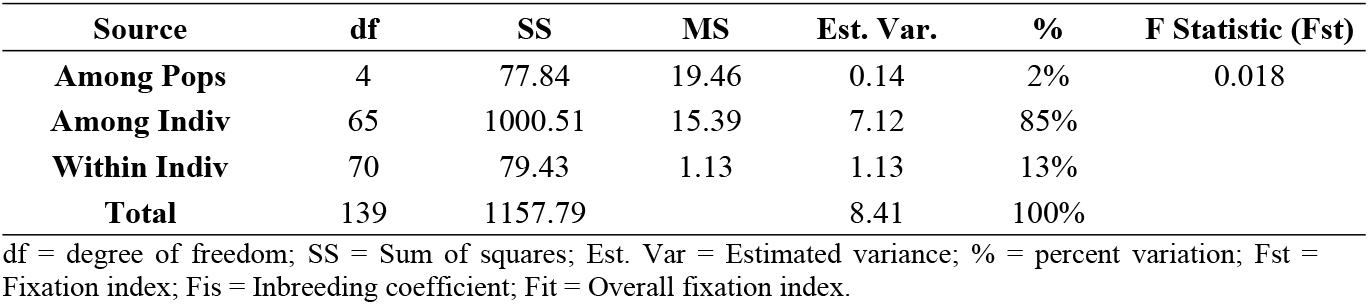
Genetic clusters and member of genotypes observed from population structure analysis of 70 cowpea genotypes.

**Fig 1.**
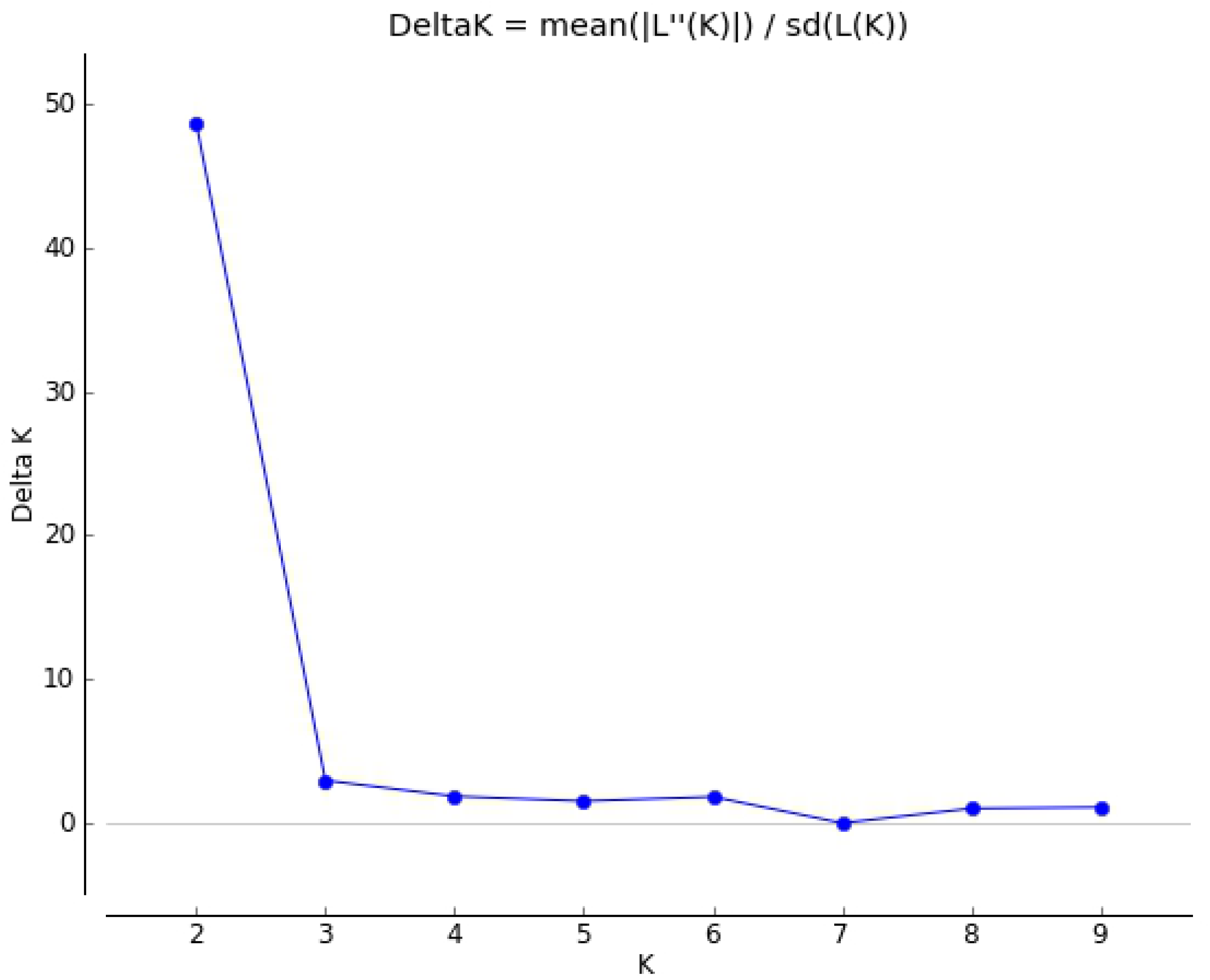
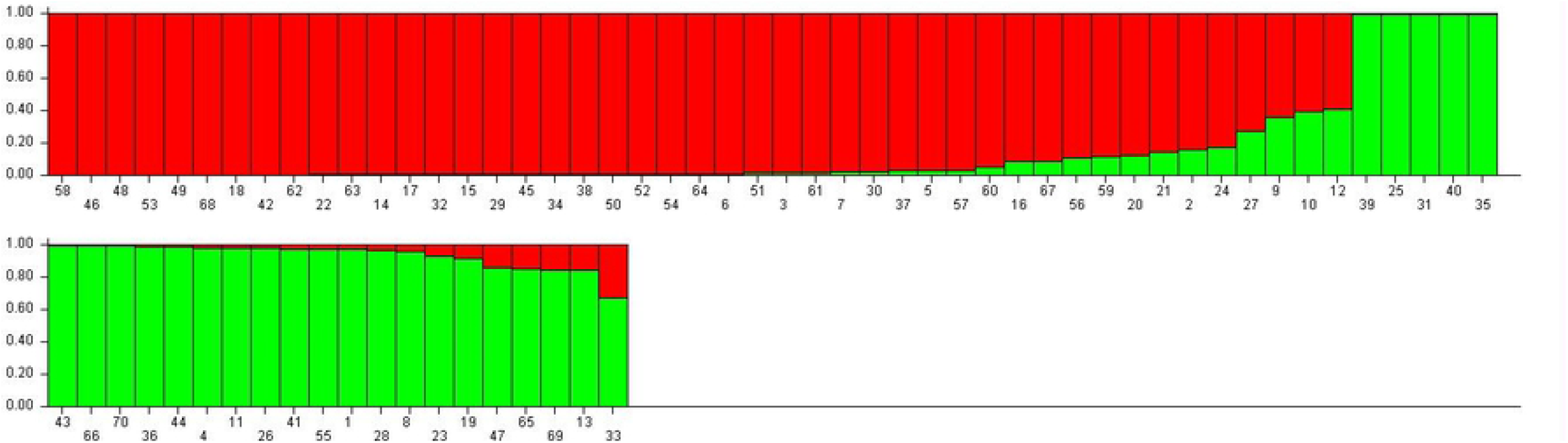
Population structure of 70 accessions based on 28 SSR markers (K = 2) and graph of estimated membership fraction for K = 2. The maximum of ad hoc measure ΔK determined by structure harvester was found to be K = 2, which indicated that the five populations could be grouped into two subgroups

**Fig 2.**
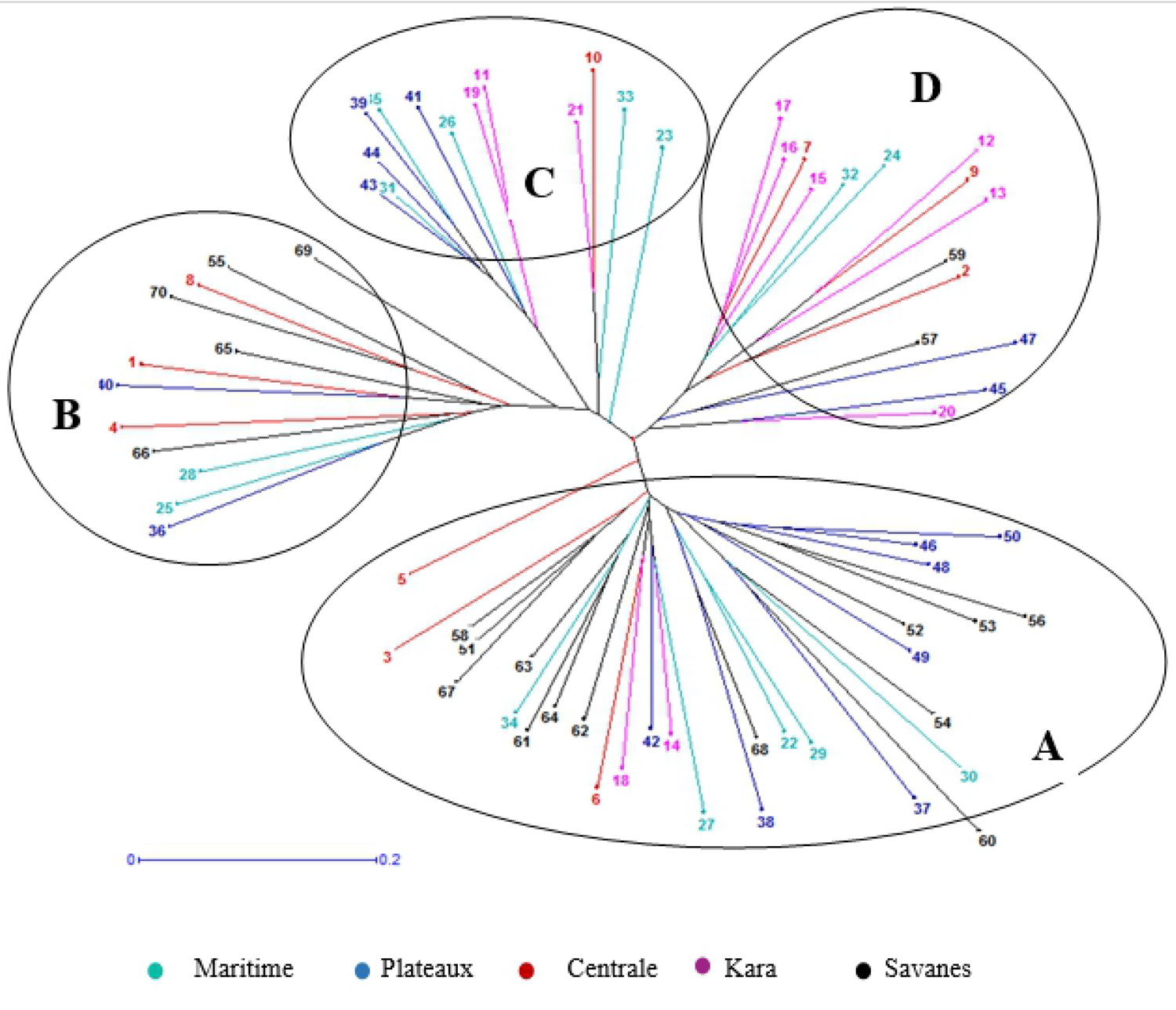
Phylogenetic tree among 70 cowpea accessions studied revealed by neighboring joining analysis.

Specific FST values were calculated for each population using STRUCTURE software. The results were respectively 0.08 for subpopulation 1 and 0.15 for subpopulation 2, with an average of 0.12 indicating a relatively low level of population structure. The average distances (i.e., expected heterozygosity) between the individuals in the same cluster were 0.53 for subpopulation 1 and 0.54 for subpopulation 2 (Table 7).

### AMOVA and Phylogenetic analysis

AMOVA was performed using the matrix of distances for genetic differentiation. The results of AMOVA revealed that the majority of variance occurred within individuals and accounted for 85% among individuals within regions of the total variation, whereas 2% and 13% of the variation was attributed to differences between population and within individuals. The results indicated that the diversity within regions (intra-regional diversity) was far greater than the diversity between regions (inter-regional diversity), and the low Fst value (0. 018) indicated a low level of dedifferentiation among regions (Table 8).

The accessions studied were clustered into four main groups. A. B. C and D based on genetic dissimilarity using the neighbour-joining method in DARwin 5.0 (Figure 3). Like the results of the structure analysis, there was no group made up exclusively of accessions from the same region. Cluster A was the largest group and contained 30 accessions. It gathered 13 accessions from the Savannah Region (43.33%), seven accessions from the Plateaux Region, five accessions from the Maritime region, three accessions from the Centrale region and two accessions from the Kara region. Group B gathered 12 accessions but did not contain any accessions from Kara Region. Group 3 contained 13 accessions from four geographical regions, with accessions from the Savannah region being absent. Group 4 contained 15 accessions, with 40% being from the Kara region with the remaining ones from the other four regions.

## Discussion

### Variability in SSR markers

An efficient evaluation of genetic resources can help reduce redundancies and build a core collection, and a core collection can be screened to identify traits of interest. Molecular markers are powerful tools for elucidating variations and relationships within and between cowpea germplasm populations. Among the genetic markers, SSRs are successfully applied in various breeding programs to study genetic diversity because of their multi-allelic nature, their level of polymorphism and the ease of their use [19,9,34,35]. Previous studies have shown that SSRs are efficient markers for genetic diversity, population structure and QTL studies using cowpea germplasm [33,36].

This study revealed a number of allele ranging from 2 to 14 allele per locus, which appeared to be relatively low compared to the value of 2 to 15 alleles per locus obtained by Sarr et al. [37] using 15 SSR markers to screen 671 cultivated cowpea from Senegal, the value of 1 to 16 obtained by Badiane et al. (2012) by screening 22 local cowpea cultivars and inbred lines collected throughout Senegal using 44 SSR markers or the value of 2 to 17 obtained by Ali et al. [38] using 16 SSR to screen 252 cowpea accessions from Sudanese germplasm. However, the range of alleles detected by loci reported in this study is wider than those reported by other studies on cowpea germplasm diversity in Senegal (1-9), Ghana (1-6), Burkina Faso (5-12), and Nigeria (2-5) [2,7,9,39]. Given the relatively lower number of alleles per loci reported by the latter cited studies, one might think that the cowpea germplasm they assessed was less diversified than the one used in our study. However, the observed difference might be explained by the difference in the number of accessions screened and the number of markers used. Indeed, Lacape et al. [40] reported that the number of amplified alleles per locus depends on the selected markers and the type of germplasm.

The estimated average PIC value (0.67) recorded in the current study was similar to the value (0.68) reported by Ogunkanmi et al. [41], higher than the values reported by Asare et al. [9]; Badiane et al. [26] and Ali et al. [38] who have respectively reported average PIC values of 0.38, 0.23, and 0.56. Therefore, SSR markers used in this study confirmed an interesting genetic diversity in the Togolese cowpea germplasm.

The average gene diversity expressed by the expected heterozygosity (He), which is a measure of genetic diversity observed in the present study (0.54), was higher than the value (0.488) reported by Ali et al. [38] for Sudanese cowpea germplasm and the value of 0.135 reported by Mafakheri et al. [42] in a study of 32 cowpea genotypes collected from different countries. However, the observed heterozygosity (0.073) revealed by the current study is very low and can be explained by the selection pressure exerted by farmers that might have reduced the polymorphism level of the germplasm.

The population structure analysis based on STRUCTURE revealed the presence of two subpopulations among the 70 cowpea accessions collected from the five regions of Togo, while Sarr et al. [37] and Xiong et al. [43] reported three populations when they respectively studied the genetic structure of 671 cultivated cowpea accessions from Senegal and the population structure of 768 cultivated cowpea genotypes from the USDA GRIN cowpea collection, originally collected mainly from around the world.

In the present study, the genetic variation components confirmed fair genetic diversity among individuals within regions (85%) than among regions (2%). The current study agrees with the findings of Sarr et al. (2020), who also reported a higher percentage variation among individuals within regions (75%). However, the percentage of variation attributed to differences between populations obtained in their study is higher than the 2% obtained in the current study. As already suggested by Sarr et al. [37], the high intra-regional diversity could be linked to the presence of many different accessions in each region. While the low genetic diversity between regions could be partly explained by the distribution of the same cowpea seed (same accessions are found everywhere) in all the regions through donations, seed companies, or agricultural extension services. Accessions from Kara seem to be genetically identical or very close to the Savannah and Centrale region, given the zero value of the differentiation indices between the Kara and Centrale regions. The observed similarity can be explained by the proximity of Kara Region to the two other regions. In fact, Kara Region is located in-between those two regions.

The value of Fst was observed to be 0.018, indicating little differentiation among populations. The fixation index (Fst) obtained in the current study was much lower than the value of 0.114 obtained for the Senegalese germplasm.

The dendrogram based on SSR markers revealed four groups. This indicates the existence of a high degree of genetic diversity in the germplasm evaluated in this study. Therefore, these germplasms could serve as a valuable source for the selection of diverse parents for a breeding program aiming to create new cultivars associating different traits of interest. However, in this study, the grouping was not observed according to regional basis or on the basis of maturity duration, habitat status or seed appearance.

## Conclusion

In Togo, cowpea is one of the main legume crops. However, the crop is poorly characterized. The current study provides useful information on the variability of SSR markers leading to a better understanding of the population structure and the genetic basis existing. It is the first study to address the genetic characterization of the Togolese germplasm, and it showed that the genetic structure does not depend on regions. The results obtained from this study will serve as basic information by providing options to breeders to develop, through selection and breeding, new and more productive cowpea cultivars that are adapted to changing environments. Furthermore, the collected germplasm could also be used for developing population for QTLs mapping studies in order to identify loci controlling traits with agronomic importance.

## Acknowledgments

We thank the farms of fives study areas for their sharing and their availability.

## Author Contributions

**Conceptualization:** Yao Dodzi Dagnon

**Data curation:** Yao Dodzi Dagnon, Koffi Tozo

**Formal analysis:** Yao Dodzi Dagnon, Koffi Kibalou Palanga

**Funding acquisition:** Yao Dodzi Dagnon

**Investigation:** Yao Dodzi Dagnon, Koffi Kibalou Palanga

**Software:** Yao Dodzi Dagnon, Koffi Tozo

**Supervision:** Koffi Tozo, Damigou Bammite

**Validation:** Koffi Tozo, Damigou Bammite. Visualization: Yao Dodzi Dagnon

**Writing – original draft:** Yao Dodzi Dagnon, Koffi Kibalou Palanga, Ghislain Comlan Akabassi

**Writing – review & editing:** Yao Dodzi Dagnon, Koffi Kibalou Palanga, Ghislain Comlan Akabassi

